# Mutually exclusive autism mutations point to the circadian clock and PI3K signaling pathways

**DOI:** 10.1101/653527

**Authors:** Hannah Manning, Brian J. O’Roak, Özgün Babur

## Abstract

Mutual exclusivity analysis of genomic mutations has proven useful for detecting driver alterations in cancer patient cohorts. Here we demonstrate, for the first time, that this pattern is also present among *de novo* mutations in autism spectrum disorder. We analyzed three large whole genome sequencing studies and identified mutual exclusivity patterns within the most confident set of autism-related genes, as well as in the circadian clock and PI3K/AKT signaling pathways.

## Main text

DNA sequencing studies have identified several genes whose mutations are strongly associated with autism spectrum disorder (ASD) as well as a greater number of weakly associated genes that require further investigation. While over 1,000 of these genes are catalogued and classified in the SFARI Gene database^1^, only 25 rank in the highest-confidence category (rank 1) (Fig.1a). We need better methods to distinguish functionally impactful genetic mutations from those that are innocuous. More importantly, we need methods to biologically contextualize these so-called functional mutations and to explain how and when they contribute to ASD etiology.

A fundamental approach for discovering functional genomic alterations is to focus on genomic regions with high numbers of variant calls in probands relative to background controls. However, such frequent mutations are rare in complex heterogeneous disorders such as ASD. In the cancer research field, we and others have employed an approach that is uniquely well-suited to this scenario: we simultaneously evaluate mutations within multiple genes to test whether their distribution across patients is non-random^2–4^. This approach identifies “mutually exclusive” gene groups whose members—while mutated throughout a disease cohort—are rarely co-mutated within an individual. The mutual exclusivity pattern indicates the existence of substitutable functional mutations, which often align with known biological pathways. Such a pattern is characteristic of certain subgroups of cancer-driving mutations and it contrasts with the random distribution of innocuous mutations (Fig. 1c). Here we show that mutual exclusivity also exists in ASD datasets and that it presents a novel opportunity for detecting and characterizing functional mutations that, to date, have been indistinguishable from randomly distributed background mutations.

**Figure 1.**
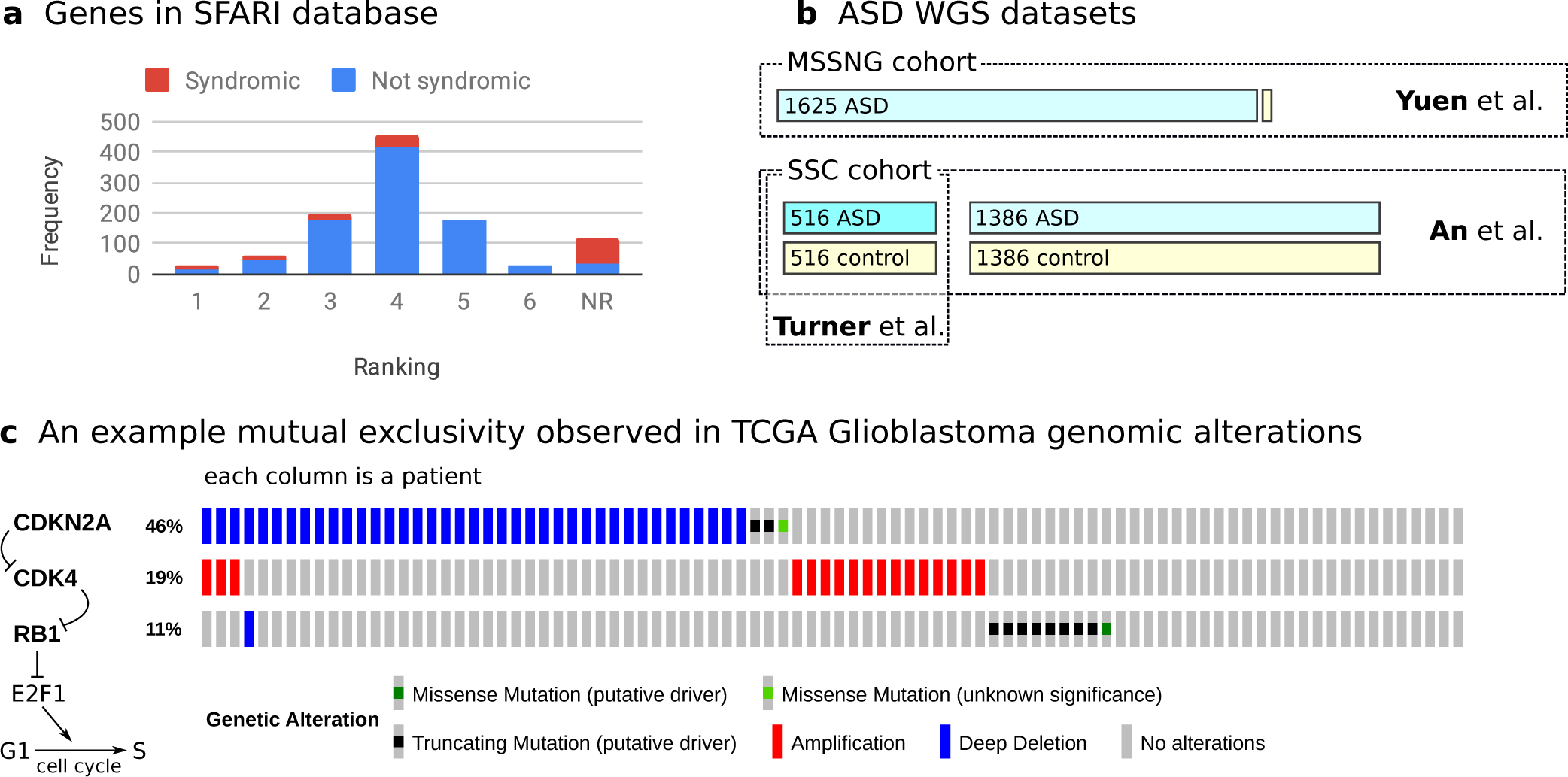
Overview of resources in ASD and the concept of mutual exclusivity in cancer. **a)** Frequency of autism genes in SFARI database grouped by their assigned ranks (lower rank indicates higher confidence). **b)** Contents of 3 whole-genome ASD datasets used in this study. **c)** An example of mutual exclusivity from the TCGA Glioblastoma cohort wherein *CDKN2A, CDK4* and *RB1* genomic alterations exhibit much less overlap than would occur randomly^2^. In this case, inhibition of either CDKN2A or RB1, or activation of CDK4 is enough to unlock the cell cycle, and a second gene alteration in this group is unnecessary for disease progression.

We use three recent whole genome sequencing ASD datasets with *de novo* mutation calls (Fig 1b): (i) a dataset released by Yuen *et al.*^5^ from the MSSNG project containing 1,625 individuals with ASD and 2 control cases, (ii) a dataset by Turner *et al*.^6^ analyzing 516 ASD probands and 516 unaffected sibling controls from Simon Simplex Collection (SSC), and (iii) a dataset published by An *et al*.^7^ analyzing 1,902 ASD probands and 1,902 unaffected sibling controls from the SSC cohort. Although all samples in the Turner dataset were re-analyzed within the An dataset, it should be noted that the Turner study’s probands were specifically selected for their lack of likely gene-disrupting (LGD) *de novo* mutations or large copy number variants (CNVs).

For proof of principle, we first test whether mutations of SFARI genes are distributed in a mutually exclusive fashion—as these genes are the most likely to bear functional mutations. To do so, we transform each dataset into a gene by sample mutation matrix and test for mutual exclusivity of the high-confidence SFARI genes with varying confidence thresholds (see Methods). We determine that the most confident SFARI gene set (rank 1) has mutually exclusive mutations in both the Yuen dataset (*p* = 0.0251) and in the combined Yuen and An dataset (*p* = 0.0108). This significance diminishes as we expand the gene set with less-confident tiers (Fig. 2a, Suppl. Tab. 1a-e). Notably, the vast majority (91%) of the gene-associated *de novo* mutations in these datasets are intronic. To understand if the observed mutual exclusivity is driven by coding mutations, we perform the same analysis on the combined Yuen and An datasets (i) using only intronic mutations, and (ii) excluding intronic mutations. While the intron-only test produces a nearly significant result for the SFARI rank 1 gene set (*p* = 0.0553, Suppl. Tab. 1f), excluding intron mutations results in only one overlap but insignificant results (*p* = 0.6421, Suppl. Tab. 1g). We conclude that, while intronic mutations may contribute to the observed mutual exclusivity, we lack the abundance of coding mutations necessary to confirm or deny their contribution.

**Figure 2.**
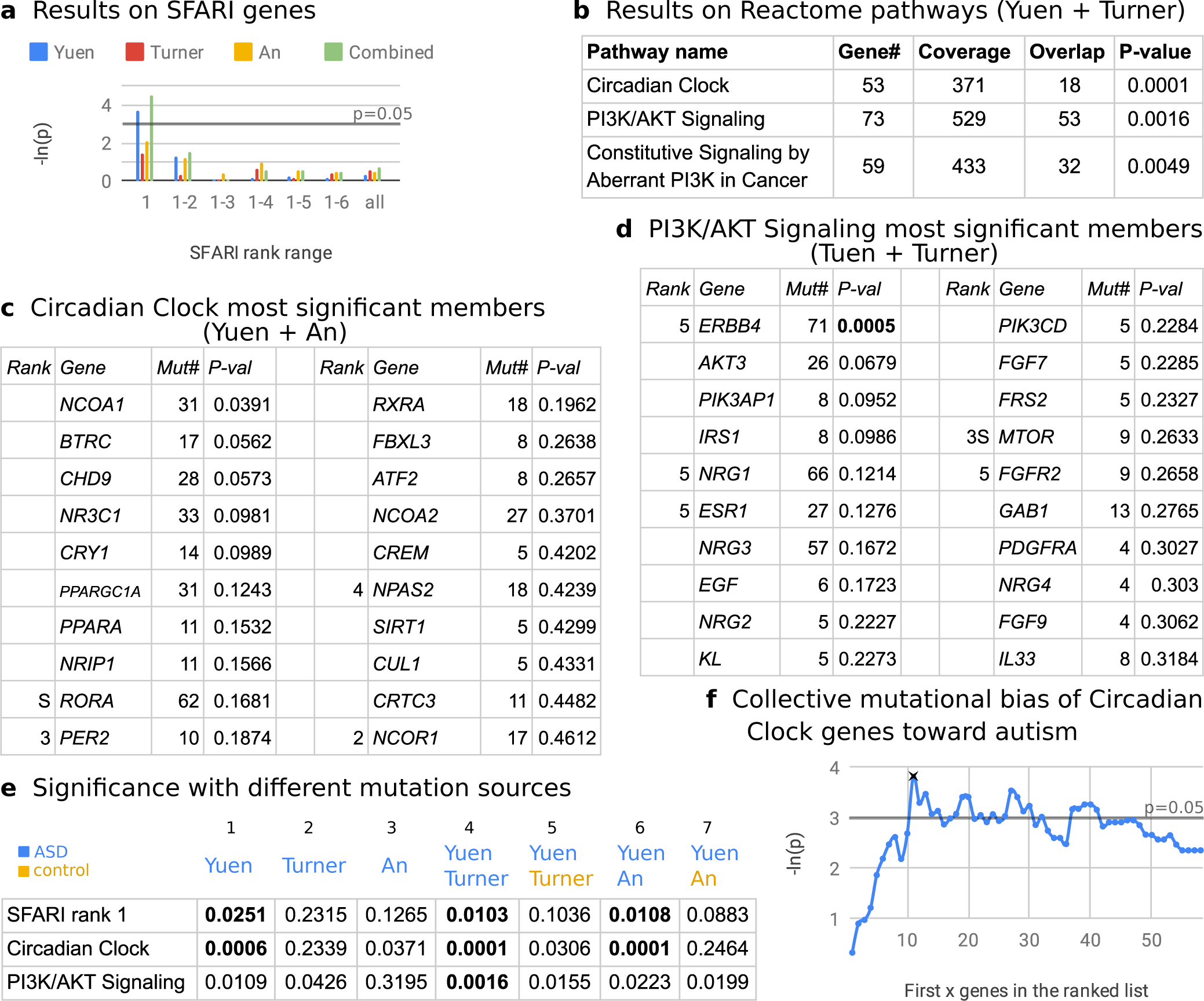
Results and validation. **a)** Mutual exclusivity test results on SFARI gene groups. The most confident group has mutually exclusive alterations. “Combined” refers to the merged Yuen and An dataset. **b)** Significant results on Reactome pathways using merged Yuen and Turner dataset. **c)** The top 20 member genes of the Circadian Clock pathway, ordered by the significance of their individual contribution to the mutual exclusivity pattern observed in the merged Yuen and An dataset, are listed. The “Rank” column shows the classification of the gene in the SFARI Gene database (empty for non-SFARI genes). The “Mut#” column shows the number of samples in which the gene is mutated at least once. **d)** The most significant 20 members of the PI3K/AKT signaling pathway in the merged Yuen and Turner dataset, arranged as in **c. e)** To verify the source of the mutual exclusivity signal, we repeat our analyses on other combinations of the WGS datasets. The first 3 columns use datasets individually while the remaining columns use combinations of the Yuen dataset with ASD or control samples from the Turner or An datasets. P-values considered significant given FDR cutoff are printed in bold. **f)** Genes in the Circadian Clock pathway are more likely to be mutated in ASD relative to the unaffected sibling controls. First x genes used are along the x axis, where the genes are ordered by the significance of their individual contribution to mutual exclusivity. Here, mutual exclusivity was tested in the Yuen dataset and mutational bias toward Autism was assessed in the An dataset.

Next, we test biological pathways from the Reactome knowledgebase^8^ to identify processes harboring mutually exclusive mutations. Reactome contains 1,576 pathways of various sizes. While exhaustively testing every pathway would not produce significant results, limiting the search space to the most mutated 50 pathways identifies the Circadian Clock pathway within the Yuen dataset (*p* = 0.0006, *FDR* = 0.03, Suppl. Tab. 2), and we find that the An dataset independently validates this result (*p* = 0.0371, Suppl. Tab. 3). To identify more pathways, we again look to the combined Yuen and An datasets, but here we find only the Circadian Clock pathway (Suppl. Tab. 4). Interestingly, using the combined Yuen and Turner dataset identifies the Circadian Clock as well as two additional pathways—both of which are related to PI3K signaling (*FDR* = 0.0817, Fig. 2b,c,d, Suppl. Tab. 5). It is possible that the sample selection criteria in the Turner dataset has, for reasons not intended nor yet understood, enriched it for functional non-coding mutations in PI3K signaling.

In the combined Yuen and An dataset, the Circadian Clock pathway has 53 mutated member genes—15 of which are catalogued by SFARI (although none of them are ranked 1, Fig. 2c, Suppl. Tab. 6, Suppl. Fig. 1). None of the member gene contributions to the observed exclusivity are individually significant. In the combined Yuen and Turner dataset, the PI3K/AKT Signaling pathway has 73 mutated member genes—11 of which are catalogued by SFARI (again, none of them are ranked 1, Fig. 2d, Suppl. Tab. 7, Suppl. Fig. 2). For this pathway, the member gene *ERBB4*, which is of low-confidence among SFARI genes (rank 5), has a significant contribution to the mutual exclusivity pattern (*p* = 0.0005, *FDR* = 0.0365). These 2 significant pathways are totally disjoint in terms of their mutated members, while the other PI3K pathway—Constitutive Signaling by Aberrant PI3K in Cancer—is a subset of the second pathway.

We perform three additional analyses to further validate our findings. First, we interrogate the reason for the observed increase in PI3K/AKT Signaling pathway mutual exclusivity brought about by replacing the An dataset with the smaller Turner dataset. Specifically, we ask whether Turner *et al.*’s unique sample selection is related to this difference rather than it being entirely attributable to general differences in data generation and processing. To test this, we calculate mutual exclusivity by using the combined Yuen and An dataset, but this time limiting the An samples to those 516 ASD probands that overlap with Turner samples. We find that, while p-values slightly deteriorate, the Circadian Clock and PI3K/AKT Signaling pathways are still significant in these results (*p* = 0.0001 and *p* = 0.0028, respectively, *FDR* = 0.07, Suppl. Tab. 8), so we conclude that sample selection has a major role in the effect, and it cannot be solely explained by data processing differences. Second, we assess the results for association with the ASD phenotype by replacing the ASD individuals with their neurotypical siblings from the Turner and An datasets (Fig. 2e, compare columns 4 versus 5 and 6 versus 7, Suppl. Tab. 1h,1i,9,10). Doing so deteriorates the p-values such that we have no significant results with either combination of datasets. We conclude that the observed mutual exclusivity is associated with the ASD phenotype. Third, we test whether the genes in the Circadian Clock pathway are more likely to be mutated in ASD than in unaffected controls. A binomial test on the An dataset using all Circadian Clock pathway members does not yield a significant result (*p* = 0.0947). However, if we sort the pathway members by their individual mutual exclusivity contributions (Suppl. Tab. 11), we find that the first 11 genes, taken together, are significantly biased toward mutation in ASD in the An dataset (*p* = 0.0218, Fig. 2f). In this analysis, we use the Yuen dataset alone for generating individual mutual exclusivity scores so that bias calculations and mutual exclusivity calculations are done on independent datasets, thereby preventing possible confounding effects.

PI3K signaling is already popularly studied in autism, as abnormalities in this pathway have been shown to promote the ASD phenotype in multiple reports^9, 10^. The circadian clock pathway, however, is relatively less studied at the genetic level, despite the well-documented prevalence of circadian rhythm disruptions within individuals with ASD^11^, and higher polymorphism of circadian clock related genes in ASD^12^. While it is still unclear if circadian rhythm abnormalities contribute to or are a by-product of ASD, our results bolster the former hypothesis and nominate the circadian clock as a key process whose alteration promotes the ASD phenotype. Here we demonstrate that mutual exclusivity analysis of *de novo* mutations is a powerful methodology for identification and characterization of functional mutations in ASD, and that it will play an important role in future research on ASD and other complex, multi-factorial disorders of unclear origins.

## Methods

### Resources

We downloaded the SFARI Gene database from https://gene.sfari.org//wp-content/themes/sfari-gene/utilities/download-csv.php?api-endpoint=genes on March 29, 2019, and provide this copy in the supplementary data.

We downloaded Reactome pathways as gene sets from the Pathway Commons^13^ database at https://www.pathwaycommons.org/archives/PC2/v11/PathwayCommons11.reactome.hgnc.gmt.gz on March 29, 2019.

We downloaded the Yuen and Turner datasets from denovo-db^14^ version 1.6.1 at http://denovo-db.gs.washington.edu/denovo-db.non-ssc-samples.variants.tsv.gz and http://denovo-db.gs.washington.edu/denovo-db.ssc-samples.variants.tsv.gz on March 26, 2019. We limited the Yuen dataset to the samples listed in their supplementary table 3 and ignored mutation calls coming from other tables that are not derived from whole genome sequencing. We downloaded the An dataset from their supplementary table 2. For each of the datasets and in every analysis, we ignored multiple mutations of the same gene in the same sample (i.e. we only considered whether the gene is mutated at least once; additional mutations on the gene in the same sample were disregarded).

### Detection of mutual exclusivity

Our mutual exclusivity detection approach is a hybrid of two previously published methods: MEMo^3^ and Mutex^2^. For a given mutation matrix and a gene set of interest, we first calculate that gene set’s “overlap” in the matrix (Fig. 3). Within the gene set, we define each sample’s overlap as one less than the number of mutated genes from the gene set in that sample (or 0 if none are present). Summation of this number across all samples produces the overlap value for the gene set within the matrix. Then, to produce a p-value for mutual exclusivity, we generate a null background distribution by shuffling the entire mutation matrix 10,000 times, preserving the number of mutations on each gene and on each sample in each shuffle, and test whether this produces the same or reduced overlap for the gene set. In addition to computing the significance for the gene set, we seek to identify the contribution of individual members. We calculate a p-value for each member gene based on its overlap with all other members collectively, using the same shuffled matrices as background.

**Figure 3.**
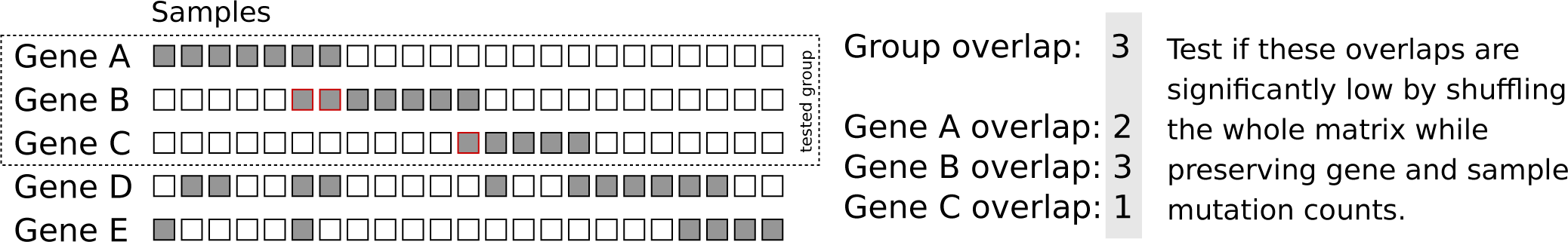
Demonstration of the mutual exclusivity detection method on a hypothetical dataset where a group—composed of genes A, B and C—is evaluated. Dark rectangles indicate that the gene is mutated in the corresponding sample (i.e. column). Red box borders highlight examples of mutations that exhibit “overlap”, i.e. are co-mutated with another gene from the group of interest.

To generate the null background distribution, we employ a degree-preserving randomization method previously applied in MEMo and originally adapted from the switching algorithm described in Milo *et al.*^15^. The algorithm preserves gene and sample mutation counts in the matrix during shuffling, as described below:

Define the mutation matrix as a bipartite graph from *G* (genes) to *S* (samples). Each edge is represented with (*g*_*i*_→*s*_*j*_)where *g*_*i*_∈*G* and *s* _*j*_∈*S. E* is the total number of edges, and *Q* is a constant. For each of our 10,000 shuffle iterations:

~~~
Do ***Q*** times
  Do ***E*** times
    Select two existing random edges (*g*_*i*_→*s*_*j*_) and (*g*_*k*_→*s*_*l*_)
    If there exist no such edges (*g*_*i*_→*s*_*l*_) and (*g*_*k*_→*s*_*j*_)
         Then replace (*g*_*i*_→*s*_*j*_) and (*g*_*k*_→*s*_*l*_) with (*g*_*i*_→*s*_*l*_) and (*g*_*k*_→*s*_*j*_)
~~~

Although there is no theoretical optimal value for *Q*, Milo *et al.* demonstrate that *Q* = 100 is a practical value for a broad variety of graphs. We follow this precedent and set *Q* to 100.

We extend MEMo’s group score with the idea of calculating individual gene scores—an idea we previously used in the Mutex approach—which enabled us to detect *ERBB4* individually and to generate ranked gene lists from the resulting gene sets. We use the Benjamini-Hochberg procedure for false discovery rate (FDR) estimation for testing the most-mutated 50 Reactome pathways. We considered 0.1 as FDR threshold for significance. In the Supplementary Tables 2-5,8-10, a pathway’s mutation count is equal to the sum of the “Coverage” and “Overlap” columns.

### Calculation of mutational bias toward autism

In the An dataset, there are a total of 73,624 mutations in probands and 72,576 mutations in the control samples. For a randomly selected mutation, the probability of belonging to a control sample is:

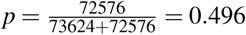

Using this probability, we define a Binomial distribution *B*(*n, p*) where *n* is the total number of mutations for a given gene or a group of genes. On the Binomial distribution, we calculate a one-sided p-value for the number of mutations in the control samples to be less than or equal to the observed value.

### Software

We have implemented the method in Java and made it available via GitHub (https://github.com/PathwayAndDataAnalysis/mutex-de-novo). We also provide a Snakemake^16^ pipeline for automated reproduction of all results reported in this manuscript, which can be run sequentially on a single machine or in parallel across a distributed system, according to the user’s resources.

## Acknowledgements

We are grateful to all of the families at the participating Simons Simplex Collection (SSC) sites, as well as the principal investigators (A. Beaudet, R. Bernier, J. Constantino, E. Cook, E. Fombonne, D. Geschwind, R. Goin-Kochel, E. Hanson, D. Grice, A. Klin, D. Ledbetter, C. Lord, C. Martin, D. Martin, R. Maxim, J. Miles, O. Ousley, K. Pelphrey, B. Peterson, J. Piggot, C. Saulnier, M. State, W. Stone, J. Sutcliffe, C. Walsh, Z. Warren, E. Wijsman). We appreciate obtaining access to data on SFARI Base. Approved researchers can obtain the SSC population dataset described in this study (SSC Whole-genome) by applying at https://base.sfari.org.

This work was supported by internal funds (B.J.O.). B.J.O. is a Sloan Research Fellow in Neurosciences (Alfred P. Sloan Foundation, FG-2015-65608 to B.J.O.) and Klingenstein-Simons Neuroscience Fellow (Esther A. & Joseph Klingenstein Fund, Simons Foundation).

## Author contributions statement

O.B. conceived the study, developed the method and the software in Java, and analyzed the datasets. B.J.O provided guidance on availability and interpretation of the genomic profiles in ASD. H.M. developed the Snakemake pipeline for reproducibility of the study and verified the results through re-runs of all analyses. All authors contributed to the writing of the manuscript and critical review of the content.

## References

1. Abrahams, B. S. et al. Sfari gene 2.0: a community-driven knowledgebase for the autism spectrum disorders (asds). Mol. autism 4, 36 (2013).

2. Babur, Ö. et al. Systematic identification of cancer driving signaling pathways based on mutual exclusivity of genomic alterations. Genome biology 16, 45 (2015).

3. Ciriello, G., Cerami, E., Sander, C. & Schultz, N. Mutual exclusivity analysis identifies oncogenic network modules. Genome research 22, 398–406 (2012).

4. Miller, C. A., Settle, S. H., Sulman, E. P., Aldape, K. D. & Milosavljevic, A. Discovering functional modules by identifying recurrent and mutually exclusive mutational patterns in tumors. BMC medical genomics 4, 34 (2011).

5. Yuen, R. K. C. et al. Whole genome sequencing resource identifies 18 new candidate genes for autism spectrum disorder. Nat. Neurosci. 20, 602 (2017).

6. Turner, T. N. et al. Genomic patterns of de novo mutation in simplex autism. Cell 171, 710–722 (2017).

7. An, J. et al. Genome-wide de novo risk score implicates promoter variation in autism spectrum disorder. Science 362, eaat6576 (2018).

8. Fabregat, A. et al. The reactome pathway knowledgebase. Nucleic Acids Res. (2018).

9. Chen, J., Alberts, I. & Li, X. Dysregulation of the igf-i/pi3k/akt/mtor signaling pathway in autism spectrum disorders. Int. J. Dev. Neurosci. 35, 35–41 (2014).

10. Kit, S. Y. et al. Identification of mutations in the pi3k-akt-mtor signalling pathway in patients with macrocephaly and developmental delay and/or autism. Mol. autism 8, 66 (2017).

11. Geoffray, M. M., Nicolas, A., Speranza, M. & Georgieff, N. Are circadian rhythms new pathways to understand autism spectrum disorder? J. Physiol. 110, 434–438 (2016).

12. Yang, Z. et al. Circadian-relevant genes are highly polymorphic in autism spectrum disorder patients. Brain Dev. 38, 91–99 (2016).

13. Cerami, E. G. et al. Pathway commons, a web resource for biological pathway data. Nucleic Acids Res. 39, D685–D690 (2011).

14. Turner, T. N. et al. denovo-db: a compendium of human de novo variants. Nucleic acids research 45, D804–D811 (2016).

15. Milo, R., Kashtan, N., Itzkovitz, S., Newman, M. E. J. & Alon, U. On the uniform generation of random graphs with prescribed degree sequences. arXiv preprint cond-mat/0312028 (2003).

16. Köster, J. & Rahmann, S. Snakemake - a scalable bioinformatics workflow engine. Bioinformatics 28, 2520–2522 (2012).

