## Supplementary Figure 1 for "Mutually exclusive autism mutations point to the circadian clock and PI3K signaling pathways"

**Supplementary Figure 1.** Circadian Clock Signaling pathway de novo gene mutation distributions. 548 samples are mutated out of 3523. Showing only mutated samples. Genes are ordered by their individual mutual exclusivity significance for this pathway.

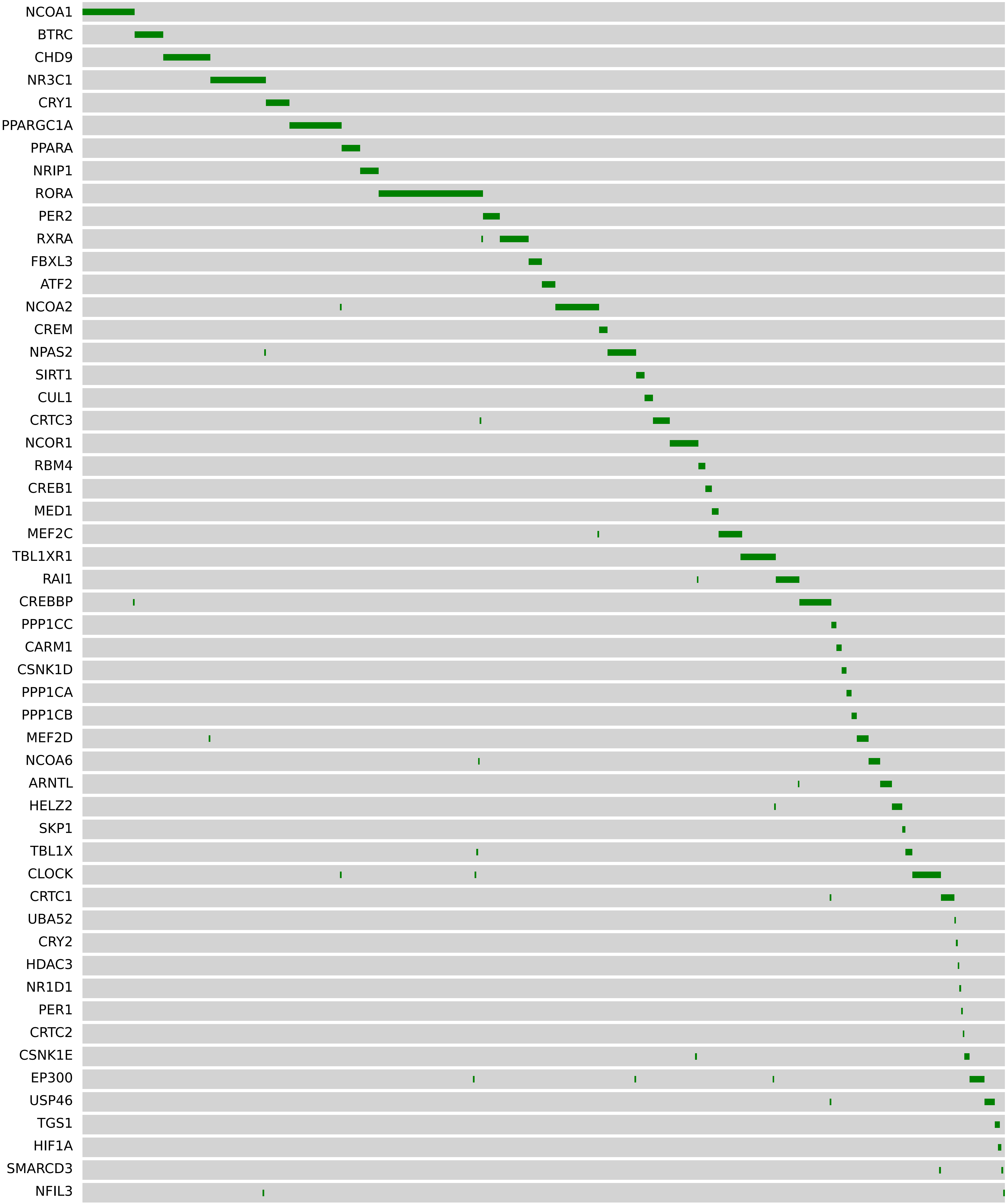
