## Supplementary Figure 2 for "Mutually exclusive autism mutations point to the circadian clock and PI3K signaling pathways"

**Supplementary Figure 2.** PI3K/AKT Signaling pathway de novo gene mutation distributions. 529 samples are mutated out of 2137. Showing only mutated samples. Genes are ordered by their individual mutual exclusivity significance for this pathway.

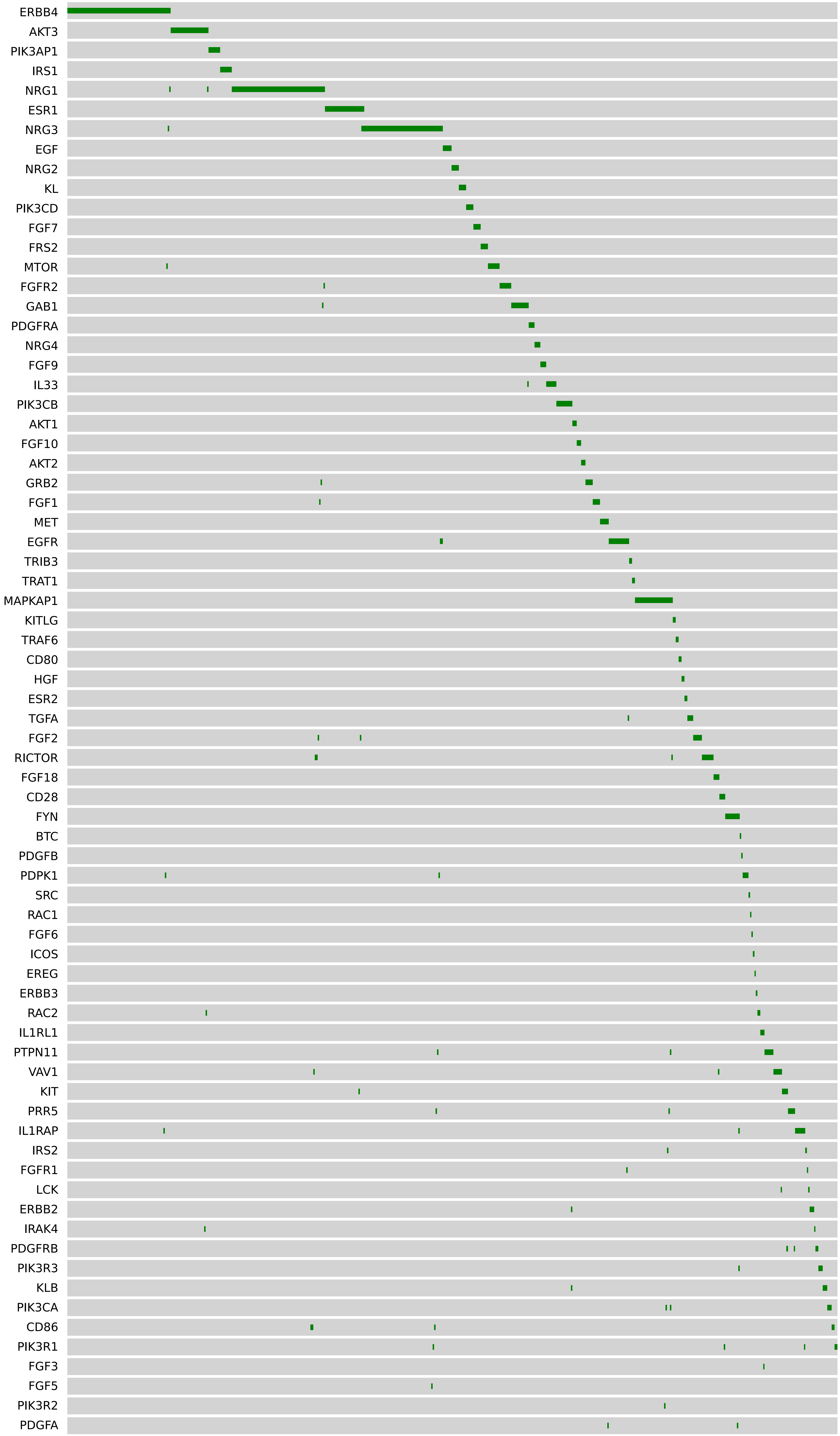
